# Distinct melanoma EV subpopulations reflect immune- and stress-induced tumor states

**DOI:** 10.64898/2026.06.16.732566

**Authors:** Michalina Sypka, Anish Ghimire, Arnau Solé Casaramona, Tanmayi Bingi, Anne-Catherine Vogt, Jie Han, Theresa Whiteside, Darien Toledo, Stephan von Gunten, Martin F. Bachmann, Mona O. Mohsen, Paul Engeroff

## Abstract

Melanoma is the deadliest form of skin cancer, and improved non-invasive approaches to monitor tumor burden and immune dynamics are needed. Although extracellular vesicles (EVs) are increasingly explored as cancer biomarkers, how distinct EV subpopulations reflect dynamic tumor states induced by immune pressure remains insufficiently understood.

Using proteomic analyses, we identified the melanoma-associated antigens gp100 (PMEL) and GPNMB in EVs derived from B16F10 melanoma cells and incorporated them into sandwich enzyme-linked immunosorbent assays (ELISAs) that capture total EVs while selectively detecting gp100⁺ and GPNMB⁺ EV subpopulations. We subsequently evaluated these EV populations in murine models of anti-tumor vaccination and in plasma samples from melanoma patients.

*In vivo,* melanoma-associated EV subpopulations increased in the serum of tumor-bearing mice and were further augmented following antigen-specific anti-tumor vaccination, whereas total CD81⁺ EVs accumulated more gradually. Notably, gp100⁺ EV levels, but not GPNMB⁺ EVs, correlated with tumor-infiltrating lymphocyte densities across multiple time points and treatment conditions. *In vitro*, TNFα/IFNγ stimulation preferentially increased total CD81⁺ EV release, whereas gp100⁺ EVs were promoted by IL-1β stimulation. In contrast, GPNMB⁺ EVs accumulated more gradually and broadly across inflammatory, hypoxic, and cytotoxic stress conditions. Together, these findings indicate that immune and stress signals differentially remodel melanoma-associated EV composition. The assay translated to humans, revealing elevated gp100⁺ EV levels in the plasma of melanoma patients compared with healthy donors, whereas GPNMB⁺ EVs identified a subset of melanoma patients.

We describe a clinically feasible approach that enables direct detection of melanoma-associated EV subpopulations from blood without prior EV isolation. Conceptually, our findings suggest that immune and stress signals dynamically shape circulating EV composition, generating distinct EV signatures that reflect tumor state.

## BACKGROUND

Melanoma is a malignant skin cancer that originates from melanocytes, specialized pigment-producing cells. Despite being responsible for only 1% of all skin cancers cases, melanoma is responsible for the majority of skin cancer–related deaths, underscoring the importance of early detection for effective treatment and survival^1,2^.

Reliable biomarkers are needed not only to assess melanoma burden, but also to predict therapy response, resistance development, and tumor behavior. In this context, liquid biopsy (LB) approaches are particularly attractive, as they enable non-invasive detection of tumor-derived material in biological fluids^3,4^.

Extracellular vesicles (EVs) are released by most cell types and are readily detectable in bodily fluids, making them well suited for LB applications^5,6^. Beyond their diagnostic potential, EVs play important roles in intercellular communication and have been increasingly implicated in cancer progression, including tumor survival, metastasis, and immune evasion^7–13^. Conversely, EVs are also being explored as therapeutic tools in oncology^14–18^. In melanoma, both preclinical models and patient studies suggest that the composition of circulating EVs changes with tumor burden, raising the possibility that melanoma-specific EV antigens could serve as biomarkers^19–21^.

However, there remains a need for rapid and cost-effective detection platforms that capture EV heterogeneity while facilitating clinical translation. Moreover, how distinct circulating EV subpopulations reflect dynamic tumor states induced by immune pressure remains poorly understood.

Here, we identify melanoma-associated EV surface antigens and establish a sandwich ELISA strategy that enables scalable, non-invasive detection of distinct melanoma-associated EV subpopulations directly from peripheral blood. Furthermore, we show that these EV subpopulations dynamically reflect immune- and stress-induced tumor states.

## RESULTS

### Purification and characterization of B16F10 melanoma-derived EVs

Extracellular vesicles (EVs) from B16F10 melanoma cells and the non-melanoma breast cancer cell line 4T1 were isolated by ultracentrifugation as described in Fig. 1A. Transmission electron microscopy (TEM) confirmed the presence of vesicles with characteristic lipid bilayer morphology in both preparations, with sizes ranging from ∼60 to 200 nm (Fig. 1B). Additional TEM images are shown in Fig. S1 and nanoparticle tracking analysis (NTA) likewise validated the EV size distribution (Fig. S2). EV identity was further confirmed by detection of the canonical EV marker CD81 by enzyme-linked immunosorbent assay (ELISA) and in flow cytometry compared to the respective isotype control (Fig. 1C-F). Treatment with 0.01% Tween disrupted vesicle integrity and abrogated CD81 detection, supporting membrane-dependent signal specificity (Fig. S2).

**Figure 1.**
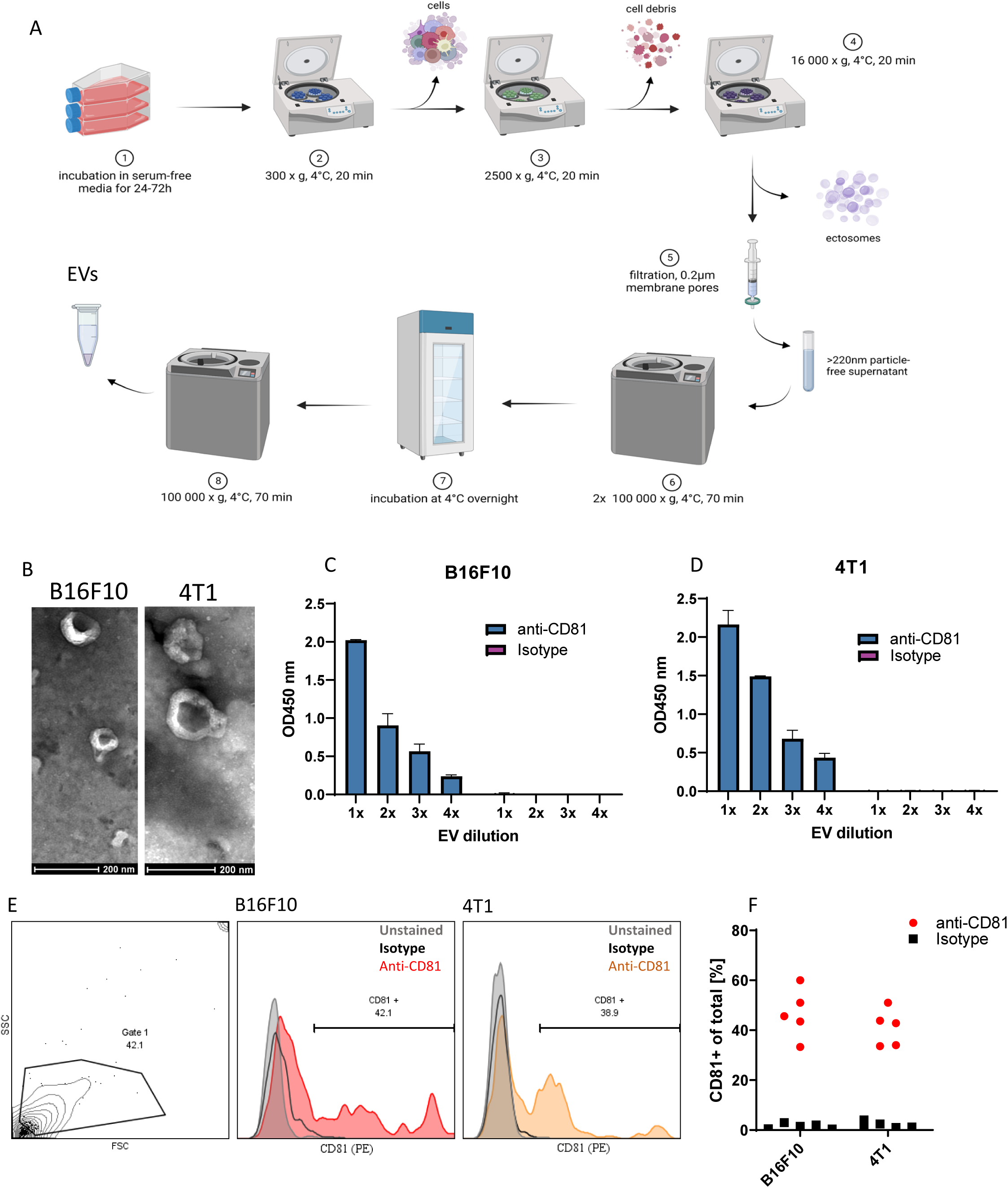
Purification and characterization of B16F10 melanoma-derived Evs. (A) Schematic representation of the ultracentrifugation workflow used for extracellular vesicle (EV) isolation. (B) Transmission electron microscopy (TEM) images of EVs isolated from B16F10 and 4T1 cell lines, showing characteristic lipid bilayer morphology and sizes. (C-D) Detection of the EV marker CD81 by ELISA in B16F10 (C) and 4T1 EV (D) preparations. (E-F) Gating strategy (E) and summarized data (F) in Flow cytometry analysis confirming CD81 expression on EVs.

Together, these data confirm successful isolation and characterization of EVs from B16F10 and 4T1 cells.

### Proteomic identification of melanoma-associated EV surface antigens

To identify melanoma-associated EV surface antigens, we performed LC–MS/MS–based proteomic profiling of B16F10 and 4T1 EVs (Fig 2). Proteomic analysis additionally confirmed EV quality by enrichment of established EV markers (CD81, CD9, CD63, TSG101, ANXA2) and absence of contaminant-associated proteins such as CYC1 and AGO1^22^ (Fig. 2A). Comparison with ExoCarta and Vesiclepedia databases confirmed substantial overlap with known EV proteomes, and several B16F10-specific proteins^23,24^ (Fig. 2B). Pathway enrichment analyses (GO, KEGG, and PANTHER) revealed that B16F10 EVs were enriched for proteins associated with melanosomes and pigment granules (Fig. 2C, S3, S4). Among the most enriched proteins found in B16F10 but not in 4T1 EVs, were the functionally related transmembrane antigens gp100 (PMEL) and GPNMB, both established melanoma-associated proteins^13,25^ (Fig. 2D, E). Consistent with the proteomic data, gp100- and GPNMB-specific antibodies selectively detected isolated B16F10 EVs but not 4T1 EVs in ELISA (Fig. 2F). Conversely, 4T1 EVs were detected via the integrin α3 antigen (Fig. S4). Based on these findings, we next developed a bi-specific sandwich ELISA that captures EVs via CD81 and detects either CD81, gp100, or GPNMB subpopulations. Indeed, all assays showed increased signals in the serum of tumor-bearing mice compared to naïve mice (Fig 2G).

**Figure 2.**
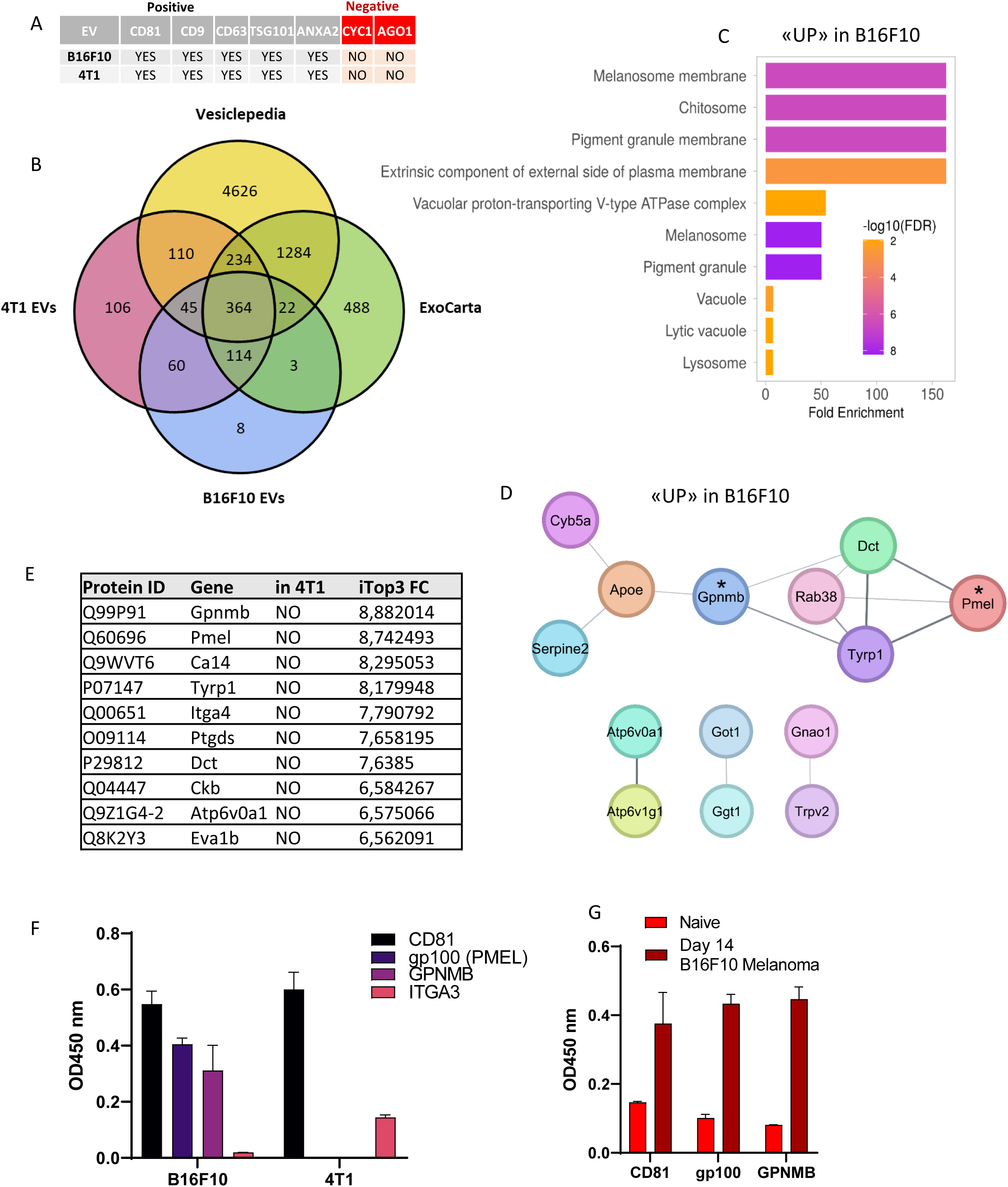
Proteomic identification of melanoma-associated EV surface antigens. (A) Proteomic validation of EV preparations by LC–MS/MS, showing enrichment of canonical EV markers and absence of contaminant-associated proteins. (B) Overlap and distinct protein composition of B16F10 and 4T1 EVs compared to ExoCarta and Vesiclepedia databases. (C) Gene ontology (GO) cellular component analysis highlighting melanosome- and pigment granule-associated pathways in B16F10 EVs. (D) Top up-regulated hits in B16F10 based on iTop3 fold changes (E) Cytoscape network representation (*top hits) of proteins in B16F10 EVs. (F-G) ELISA of gp100^+^ and GPNMB^+^ EVs in isolated EVs (F) and serum of naïve and tumor-bearing mice (G).

**Figure 3.**
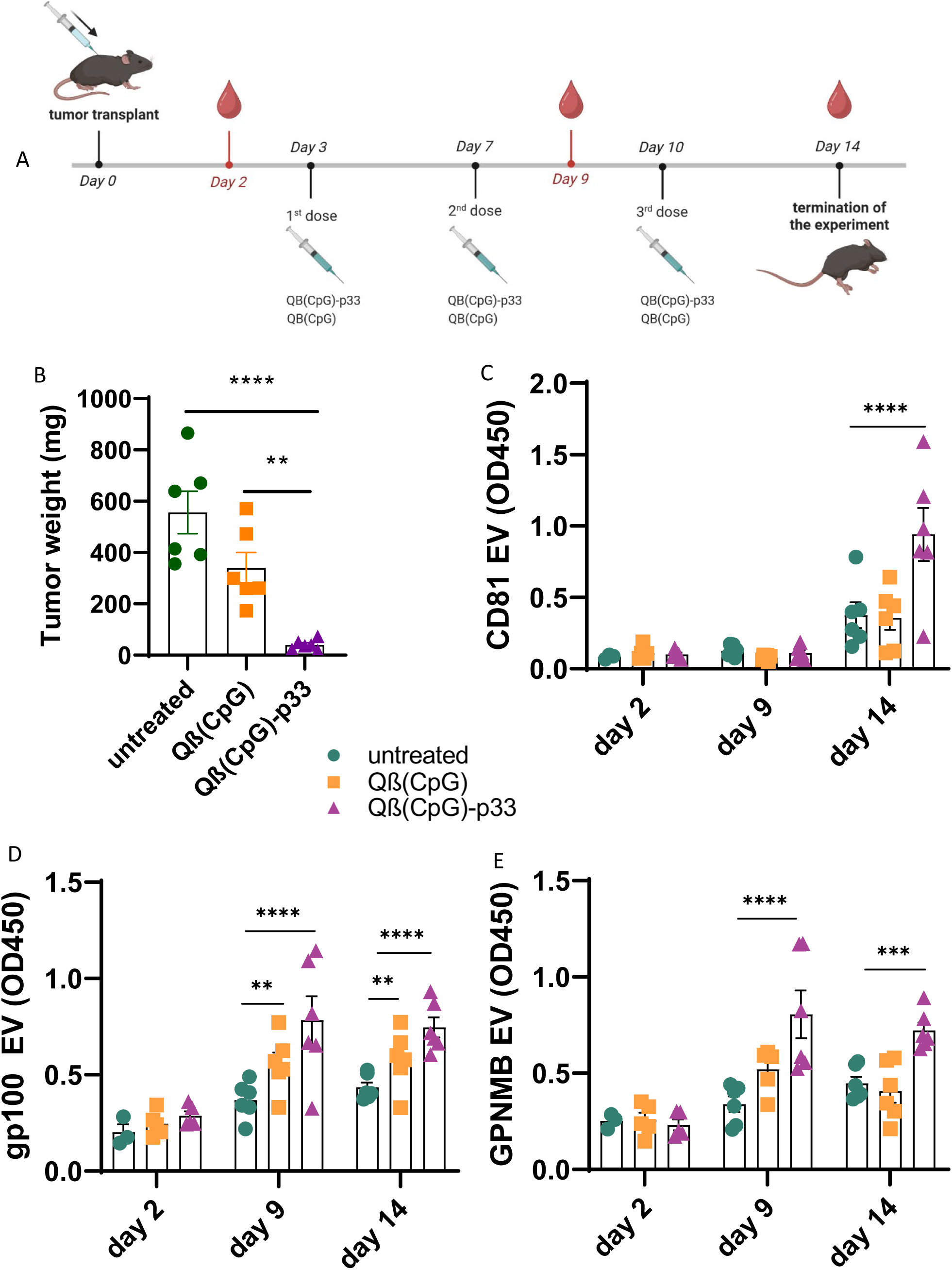
Melanoma-associated EVs increase in mice upon therapeutic vaccination. (A) Experimental design of the in vivo B16F10 melanoma model. Mice were inoculated with tumor cells on day 0 and treated with Qβ(CpG)-p33 vaccine, control Qβ(CpG) virus-like particles (VLPs), or left untreated. Vaccinations were administered on days 3, 5, and 7, and blood samples were collected on days 2, 9, and 14. (B) Tumor weight (C-D) Serum levels of CD81⁺ (C), gp100⁺ (D), and GPNMB⁺ E) EVs measured by ELISA at the indicated time points. Statistical significance was determined using two-way ANOVA with Tukey’s multiple comparisons test. All data are shown as mean ± SEM.

**Figure 4.**
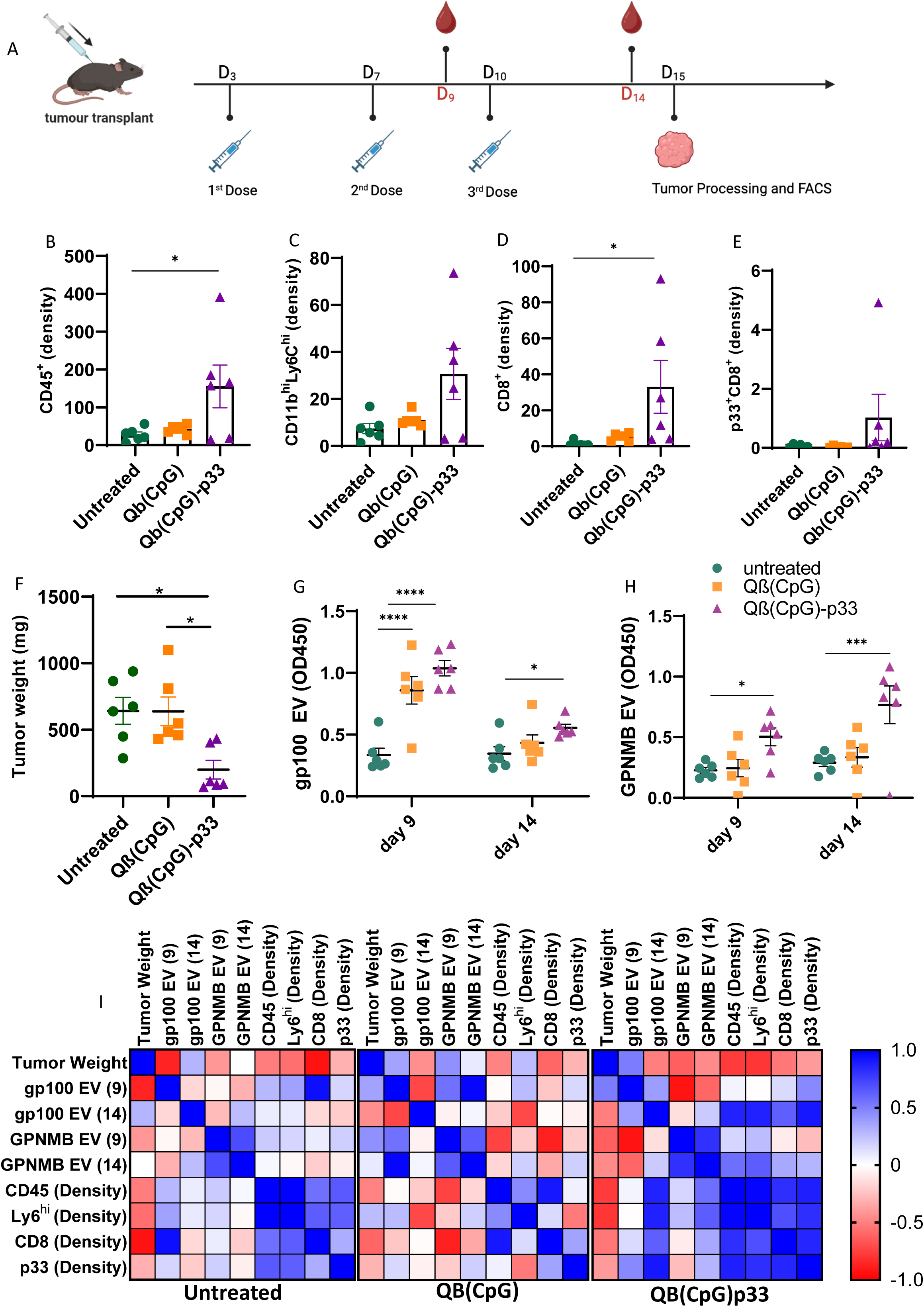
gp100⁺ EVs correlate with tumor-infiltrating immune responses. (A) Experimental overview and flow cytometry gating strategy for tumor-infiltrating lymphocytes (TILs). (B) Quantification of CD45⁺ immune cells, (C) CD11b^hi^ Ly6C^hi^ macrophages, (D) CD8⁺ T cells, (E) p33 tetramer–specific CD8⁺ T cells in tumors. (F) Tumor weight at endpoint. (G–H) Serum levels of gp100⁺ and GPNMB⁺ EVs in the corresponding animals. Statistical significance was determined using two-way ANOVA with Tukey’s multiple comparisons test. All data are shown as mean ± SEM. (I) Pearson correlation analyses between EV levels, tumor weight, and TIL densities.

Together, these findings identify gp100 and GPNMB as melanoma-associated EV surface antigens suitable for our assay system.

### Melanoma-associated EVs increase in mice upon therapeutic vaccination

We next aimed to investigate these markers longitudinally in a model of anti-melanoma vaccination. Mice were treated with a previously described virus-like particle (VLP)-based vaccine, Qβ(CpG)-p33, which induces antigen-specific anti-tumor immunity in p33-expressing B16F10 tumors^26,27^. Control groups included mice treated with non-specific Qβ(CpG) VLPs and untreated animals. Vaccination was performed subcutaneously on days 3, 5, and 7 following tumor inoculation (day 0), and blood samples were collected on days 2, 9, and 14, where mice reach humane endpoint (Fig. 3A)^26,27^.

Consistent with previous findings, Qβ(CpG)-p33 significantly reduced tumor growth compared to control-vaccinated and untreated groups (Fig. 3B, Fig S6). In this setting, serum levels of CD81⁺, gp100⁺, and GPNMB⁺ EVs increased over time, with the most pronounced changes observed in the antigen-specific treatment group (Fig. 3C-E, Fig. S5,6). Notably, gp100⁺ and GPNMB⁺ EVs increased in tumor-bearing mice as early as day 9, whereas total CD81⁺ EVs predominantly rose on day 14.

Together, these data demonstrate that EV surface antigen–based sandwich ELISAs enable dynamic monitoring of total EV versus melanoma-associated EV subpopulations directly from serum during anti-tumor immunity.

### gp100⁺ EV levels correlate with tumor-infiltrating lymphocytes

We next assessed how circulating EV markers relate to tumor-infiltrating immune responses. gp100⁺ and GPNMB⁺ EV levels were compared to tumor-infiltrating lymphocytes (TILs) at endpoint, quantified by flow cytometry (Fig. 4A, Fig. S7). Tumors were analyzed for CD45⁺ immune cells, CD11b^hi^ Ly6C^hi^ macrophages, CD8⁺ T cells, and p33 tetramer–specific CD8⁺ T cells, as previously described^26,27^ (Fig. S7).

Antigen-specific vaccination significantly increased the density of CD45⁺ TILs (Fig. 4B), and CD11b^hi^ Ly6C^hi^ macrophages showed a non-significant trend toward increased infiltration following vaccination (Fig. 4C). CD8⁺ T cells were significantly increased (Fig. 4D), whereas p33-specific CD8⁺ T cells were only increased in a subset of vaccinated tumors, consistent with heterogeneous but detectable antigen-specific responses (Fig. 4E). Tumor weights were again significantly reduced upon specific vaccination (Fig. 4F, Fig. S6). In parallel, gp100⁺ and GPNMB⁺ EV levels again increased in vaccinated animals (Fig. 4G, H).

We then performed Pearson correlation analyses between EV levels, tumor weight, and TIL densities. TIL populations positively correlated with each other, consistent with coordinated immune infiltration. Notably, in the untreated groups, gp100⁺ EV levels at day 9 inversely correlated with tumor weight (r = −0.866) and positively correlated with endpoint CD8⁺ T cell infiltration (r = 0.948).

At day 14, gp100⁺ EV levels correlated with CD45⁺ (r = 0.862), CD11b^hi^ Ly6C^hi^ macrophages (r = 0.868), CD8⁺ T cells (r = 0.625), and p33-specific CD8⁺ T cells (r = 0.913), predominantly in antigen-specific vaccinated animals but not in untreated controls (Fig. 4I, Fig. S7). In contrast, GPNMB⁺ EVs showed weaker and less consistent associations with TIL populations.

Together, these data indicate that gp100⁺ EVs associate more strongly with anti-tumor immune responses than GPNMB⁺ EVs and may function as circulating indicators of immune activation in melanoma.

### EV surface antigens reflect stimuli-dependent immune pressure

We next investigated *in vitro*, whether the identified EV surface antigens can detect stimulus-dependent changes in EV release by melanoma cells. B16F10 cells express cytokine receptors and possibly employ them to regulate metastasis or anti-tumor immunity^28,29^. Thus, B16F10 were cultured for 24 h or 48 h in the presence of TNFα/IFNγ, IL-1β. Alternatively, the cells were subjected to hypoxic conditions with CoCl2, or cytotoxic stress with doxorubicin (Fig. 5A–F).

**Figure 5.**
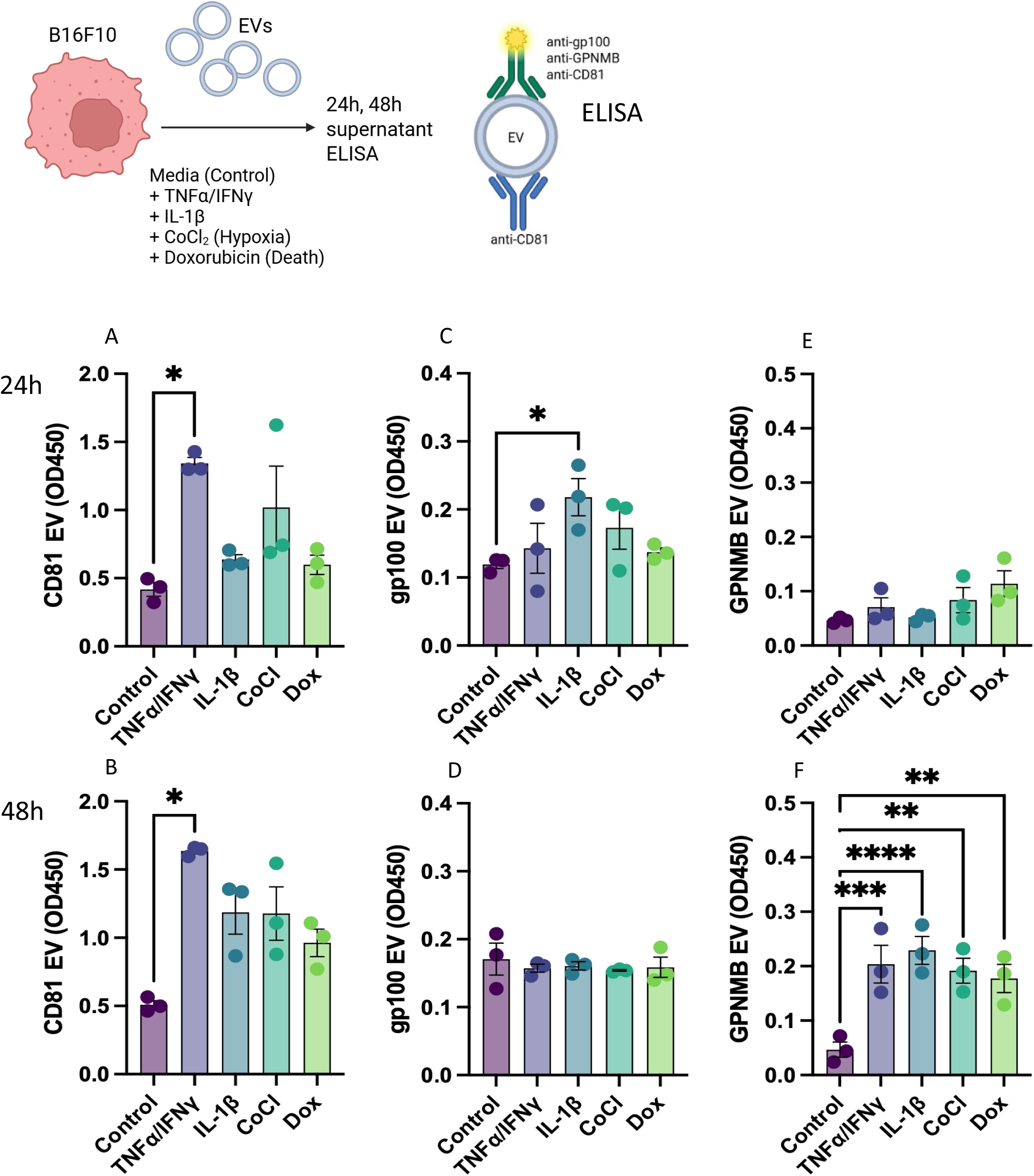
EV surface antigens reflect stimuli-dependent immune pressure. (A–F) In vitro analysis of EV release from B16F10 melanoma cells cultured under distinct stress conditions. Cells were treated with inflammatory stimuli (TNFα/IFNγ or IL-1β), hypoxic stress (CoCl₂), or cytotoxic stress (doxorubicin) for 24 h or 48 h. (A, B) Total EV release quantified by CD81⁺ ELISA. (C, D) gp100⁺ EV levels. (E, F) GPNMB⁺ EV levels. Statistical significance was determined using Kruskal–Wallis test with Dunn’s multiple comparisons test.

TNFα/IFNγ significantly increased total CD81⁺ EV release but not gp100⁺ EVs at both time points (Fig. 5A, B). In contrast, IL-1β induced a transient increase in gp100⁺ EVs at 24 h, which was not sustained at 48 h (Fig. 5C, D). GPNMB⁺ EVs showed minimal early changes but were consistently increased at 48 h across all stress conditions, cytokine stimulations, hypoxia and cytotoxic stress (Fig. 5E, F).

Together, these findings suggest that inflammatory and stress signals dynamically remodel melanoma-associated EV composition, generating distinct gp100⁺ and GPNMB⁺ EV subpopulations that reflect specific immune- and stress-induced tumor states.

### gp100⁺ EVs distinguish melanoma patients from healthy donors

The EV markers identified in the B16F10 model, particularly gp100, have previously been observed in human melanoma-derived exosomes¹³. We therefore assessed whether our ELISA-based approach could be translated to detect circulating EVs in human plasma samples.

To maximize EV capture, we adapted the assay by combining capture antibodies against CD81, CD63, and CD9 (Fig. 6A). Indeed, EVs could be readily detected from human melanoma cell line supernatants (Mel 526), both following isolation and in media supernatants (Fig. S8).

**Figure 6.**
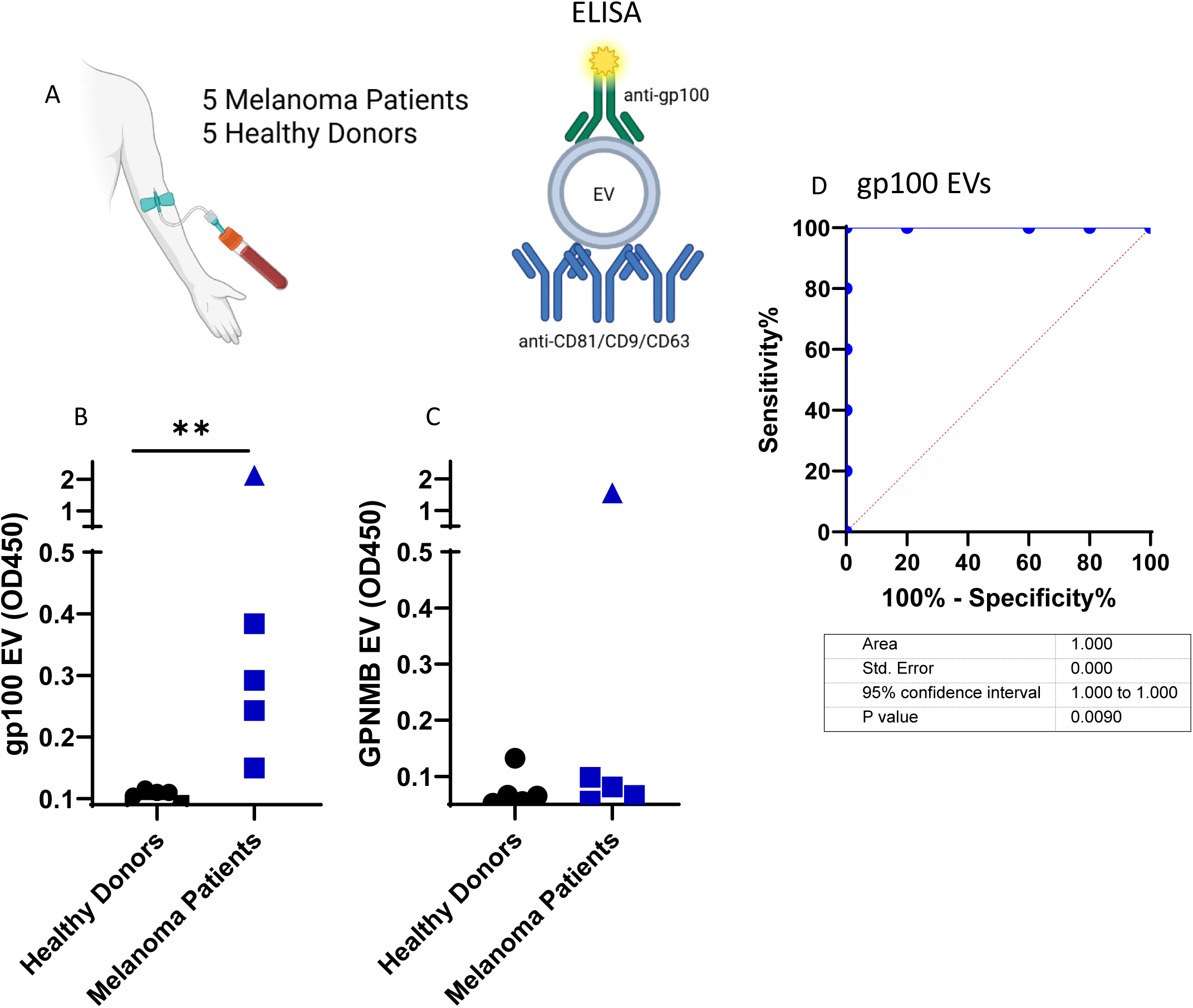
gp100⁺ EVs distinguish melanoma patients from healthy donors. (A) Schematic of the adapted EV capture strategy using a combination of anti-CD81, anti-CD63, and anti-CD9 antibodies. (B) ELISA-based detection of gp100⁺ EVs in plasma from melanoma patients and healthy donors. Statistical significance was determined using the two-tailed Mann–Whitney test. Each dot represents one individual. (C) ELISA-based detection of GPNMB⁺ EVs in the same cohort. (D) Receiver operating characteristic (ROC) curve analysis of gp100⁺ EVs.

We next measured gp100⁺ and GPNMB⁺ EVs in plasma samples from five healthy donors and five melanoma patients. Consistent with the murine data, gp100⁺ EV levels were elevated in all melanoma patients compared to healthy donors (Fig. 6B). Interestingly, one patient exhibited markedly higher gp100⁺ EV levels (OD₄₅₀ ∼2.0), whereas the remaining patients showed intermediate signals (OD₄₅₀ ∼0.2–0.4), and healthy donors were close to the limit of detection. In contrast to gp100⁺ EVs, GPNMB⁺ EV levels did not clearly distinguish patients from healthy donors. However, the single patient exhibiting very high gp100⁺ EV levels also showed strongly elevated GPNMB⁺ EVs (Fig. 6C).

These findings indicate that gp100⁺ EVs but not GPNMB⁺ EVs can discriminate melanoma patients from healthy individuals and support the potential clinical utility of EV-based ELISA assays (Fig 6D). In turn, GPNMB⁺ EVs may provide complementary information, potentially identifying patient subgroups.

## DISCUSSION

We identify the melanoma-associated EV surface antigens gp100 (PMEL) and GPNMB as markers of distinct melanoma-associated EV subpopulations that can be detected directly from peripheral blood using an ELISA-based approach. In murine models, both markers increased with tumor progression, yet their dynamics were strongly modulated by immune activation. Notably, gp100⁺ EVs correlated with TIL responses and distinguished melanoma patients from healthy donors, whereas GPNMB⁺ EVs, which were broadly induced under stress conditions *in vitro*, may reflect more general tumor stress states and inter-patient heterogeneity. A limitation of this study is the modest human sample size, which will require validation in larger and longitudinal patient cohorts.

Our findings are consistent with previous reports identifying gp100 in melanoma-derived EVs^13^. GPNMB is a well-characterized immunomodulatory protein overexpressed in melanoma and other cancers, where it has been implicated in tumor progression and immune regulation^25,30,31^. More broadly, LB approaches are increasingly adopted in oncology due to their non-invasive nature and their ability to capture dynamic tumor changes. Within this context, EVs have emerged as promising biomarkers across multiple malignancies^19–21,32^.

EV populations are increasingly recognized as highly heterogeneous with respect to their cellular origin, molecular composition, and biological function, yet to capture EV heterogeneity and resolve biologically distinct subpopulation signatures remains challenging^33–35^. Another major barrier to the clinical translation of EV-based biomarkers has been the reliance on complex and time-consuming isolation workflows. Here, we demonstrate that distinct melanoma-associated EV subpopulations can be detected directly from serum or plasma using a bi-specific ELISA strategy that combines EV capture with selective surface antigen detection. Importantly, this platform is modular and could, in principle, be adapted to different EV markers and disease contexts. Future studies will be required to define sensitivity, specificity, and clinical utility in larger patient cohorts and across therapeutic settings, as well as to determine whether this concept can be extended to additional cancer types.

An important observation of this study is that EV levels do not solely reflect tumor burden but are shaped by immune pressure. While EV signals increased with tumor growth in untreated animals, immunotherapy further enhanced EV levels despite tumor regression. Both active EV secretion and passive vesicle release from stressed or dying tumor cells have previously been described, and EVs have been implicated in tumor adaptation and immune modulation^36,37^. In addition, tumor-derived EVs can contribute to immune suppression, for example through the transfer of immune regulatory molecules. Our data support a model in which tumor cells dynamically alter EV composition in response to immune pressure, generating distinct EV signatures that reflect immune activation and stress^6,38–40^.

In summary, we identify EV surface antigens that enable non-invasive monitoring of melanoma burden and immune dynamics directly from blood. By eliminating the need for EV isolation, this ELISA-based approach provides a scalable strategy for the clinical implementation of EV-based liquid biopsies. Conceptually, our findings suggest that EV surface antigens capture stimulus-dependent tumor states and can serve as accessible reporters of tumor–immune interactions. Further studies in larger, longitudinal cohorts will be required to validate these findings and to define their utility across different therapeutic contexts.

## METHODS

### Experimental design

This study was designed to identify melanoma-associated EV surface antigens and evaluate their potential as biomarkers of tumor burden, immune activation, and stress responses. The experimental workflow combined in vitro stimulation assays, in vivo tumor models, and exploratory analyses of human samples.

In vitro experiments were performed using B16F10 melanoma cells and, where indicated, 4T1 breast cancer cells as a non-melanoma control. Cells were cultured under defined stress conditions, including inflammatory stimulation EVs were isolated from media and analyzed using ELISA-based detection of CD81⁺, gp100⁺, and GPNMB⁺ vesicles. Experiments were performed in independent biological replicates as indicated in the figure legends.

Sample sizes for animal experiments were selected in accordance with the principles of reduction (3Rs), balancing statistical power with ethical considerations, and are consistent with commonly used group sizes in preclinical immunotherapy studies. The in vivo experiments were performed at least twice independently. Investigators were not blinded to group allocation during experiments.

Human plasma samples from melanoma patients and healthy donors were analyzed to assess the translational potential of EV-based biomarker detection. These analyses were exploratory and performed on a limited cohort.

Statistical analyses, including comparisons between groups and correlation analyses, are described in the corresponding figure legends.

### Cell lines

B16F10 (mouse melanoma cells) and 4T1 (mouse breast cancer cells) were cultured in Dulbecco’s Modified Eagle’s Medium DMEM (Gibco) with 10% heat-inactivated FBS. The culture was maintained at 37°C in a humidified 5% CO2 atmosphere. In stimulation experiments, control was media, IFNy 20ng/mL + TNFa 10 ng/mL (STEMCELL), IL-1β 10 ng/mL (Thermo Fisher) CoCl₂ 150 µM (Sigma), Doxorubicin 5uM (Sigma). The human melanoma cell line Mel526 (ATCC) was used for generating human melanoma-EVs. Cells were cultured at 37 °C in 5% CO₂. Mel526 cells were maintained in RPMI-1640 supplemented with penicillin (100 U/mL), streptomycin (100 µg/mL), and 10% heat-inactivated FBS. Cells were seeded at 4 × 10⁶ cells per 150 cm² flask in 25 mL medium.

### Isolation of EVs by Ultracentrifugation

The isolation procedure for Mel526 cells was previously described^41^. For B16F10 and 4T1, cells were cultured in serum-free medium for 48 hours. Next, the culture was transferred to a falcon tube and was centrifuged at ×300 g, 4°C for 20 minutes. The pellet was discarded, and the supernatant was transferred to another falcon tube and centrifuged at ×2 500 g, 4°C for 20 minutes. After the centrifugation, the pellet was discarded and the supernatant was transferred to another falcon tube and centrifuged at ×16000 g, 4°C for 20 minutes. The supernatant was filtered through a 0.22 μm syringe filter and the solution was transferred to 13.2 mL, Open-Top Thinwall Ultra-Clear Tube, 14 x 89mm (Beckman Coulter Life Sciences) and was centrifuged at ×100 000 g, 4°C for 80 minutes. The supernatant was discarded and the tube was topped up with DPBS (Biowest). The solution was centrifuged at ×100 000 g, 4°C for 80 minutes. The supernatant was discarded; the tube was sealed with parafilm and incubated at 4°C overnight. Upon the completed incubation, the tube was filled up with DPBS and centrifuged at ×100 000 g, 4°C for 80 minutes. The supernatant was discarded; the remaining pellet was resuspended and transferred to a 1.5 mL Eppendorf tube.

### Nanoparticle Tracking Analysis

The isolated EVs were analysed via NTA using NanoSight NS300 (Malvern Instruments) featuring 405 nm blue laser and a sCMOS camera. The samples were diluted 1:100 with sterile DPBS. The camera level was set to 12, slider shutter to 1200, slider gain to 146 at 25 frames per second. Five separate measurements were conducted to assess the size and concentration of EVs.

### Quantification of EVs by Flow Cytometry

From purified EVs, 5 μl were used for counting. The EV solution of mouse origin was incubated at 4°C for 30 minutes with the detection antibody anti-mouse CD81 or with an isotype control set at the concentration 1:200 (Table 1). Flow cytometry was performed utilizing the CytoFLEX S 4L 13C (B2-R3-V4-Y4) instrument with a 96-well plate loader (Beckman Coulter Life Sciences). Analysis was carried out using CytExpert software (Beckman Coulter Life Sciences) and FLOWJO software (TreeStar Inc).

**Table 1:** Antibodies.

| Product | Conjugate | Dilution | Clone | Method | Company |
| --- | --- | --- | --- | --- | --- |
| Anti-mouse CD81 | PE | 1:200 | Eat2 | Flow Cytometry | BD Biosciences |
| Anti-mouse CD81 | Biotin | various | Eat2 | ELISA | BioLegend |
| Anti-mouse CD81 | - | $4\mu\text{g/mL}$ | Eat2 | ELISA | BioLegend |
| Anti-mouse GPNMB | - | various | Polyclonal | ELISA | ThermoFisher Scientific |
| Anti-mouse gp100 | - | various | Polyclonal | ELISA | ThermoFisher Scientific |
| Anti-rabbit IgG (H+L) | HRP | various | Polyclonal | ELISA | Nordic MUBio |
| Streptavidin | HRP | various |  | ELISA | ThermoFisher Scientific |
| Anti-hamster IgG Cocktail | HRP | various | Polyclonal | ELISA | BD Biosciences |
| Anti-human IgG | HRP | various | Polyclonal | ELISA | ThermoFisher Scientific |
| Anti-rat IgG | HRP | various | Polyclonal | ELISA | BioLegend |
| Anti-mouse IgG | HRP | 1:1000 | Polyclonal | ELISA | ThermoFisher Scientific |
| Anti-mouse IgG | FITC | 1:200 |  | Flow Cytometry | Santa Cruz Animal Health |
| Anti-mouse CD16/32 (Blocking) | - | $0.5\mu\text{g/mL}$ | 2.4G2 | Flow Cytometry | BD Biosciences |
| Anti-mouse CD3e |  | 1:400 |  | Flow Cytometry | Thermo Fisher Scientific |
| Anti-mouse CD8 | PECy7 | 1:300 | 53-6.7 | Flow Cytometry | BD Biosciences |
| Anti-mouse CD11b | APC-Cy7 | 1:300 | M1/70 | Flow Cytometry | BioLegend |
| Anti-mouse Ly6C | HK1.4 | 1:300 | APC | Flow Cytometry | BioLegend |
| Anti-mouse CD45 | BUV737 | 1:300 | 30-F11 | Flow Cytometry | Thermo Fisher Scientific |
| P33-Tetramer | PE | 1:50 |  | Flow Cytometry | MBL Life Science |
| Anti-human gp100 |  | 1:500 | MA5-101023R | ELISA | Thermo Fisher Scientific |
| Anti-human CD9 |  | 1 µg/mL | 10626D | ELISA | Thermo Fisher Scientific |
| Anti-human CD63 |  | 1 µg/mL | 10630D | ELISA | Thermo Fisher Scientific |
| Anti-human CD81 |  | 1 µg/mL | 10628D | ELISA | Thermo Fisher Scientific |

### Transmission Electron Microscopy of EVs

Physical stability and integrity of obtained EVs were visualized by transmission electron microscopy (Philips CM12 EM). For imaging, sample-grids were glow discharged and 10 μl of purified EVs were added for 30s. Grids were washed 3× with ddH2O and negatively stained with 5 μl of 5% uranyl acetate for 30s. Excess uranyle acetate was removed by pipetting, and the grids were air dried for 10 minutes. Images were taken with 84.000× and 110.000× magnification.

### Proteomic Analysis (PMSCF)

For proteomic analysis, purified EVs were then supplemented with iodoacetamide at a final concentration of 10 mM, followed by a further incubation at room temperature for 20 min. These samples were diluted to 1.6 M urea and 50 mM TrisHCl (pH 8.5) using TrisHCl (100 mM, pH 8.1), and digested with trypsin/LysC (1.5 µg of enzyme per 250 µL of diluted sample) at 37°C. After digestion, samples were acidified in 1% trifluoroacetic acid (TFA) and desalted on TARGA C18 spin columns following the manufacturer’s instruction (P/NHMMS18R, Nest Group). Desalted peptides were separated on a 75 µm × 50-cm nano-LC column packed with 2-µm C18 particles (P/N 964942, Thermofischer) using a 120 min gradient of water/acetonitrile in 0.1% formic acid. Eluted peptides were ionized using stainless-steel emitters and analyzed via “high/low”-mass accuracy data-dependent HCD MS/MS in a quadrupole-Orbitrap mass spectrometer (Fusion model, Thermofischer) in the proteomics core facility PMSCF from the Department for BioMedical Research (DBMR) at the University of Bern, Switzerland. ShinyGO 0.77 database (http://bioinformatics.sdstate.edu/go/)

### Sandwich ELISA for EVs Present in Mouse Sera

To determine the amount of EVs present in sera, ELISA Corning™ 96-Well Half-Area plates (Sigma) were coated with anti-mouse CD81 antibody at concentrations of 4 μg/mL overnight at 4°C. In initial experiments, isolated EVs were directly coated (Fig 1). For all washing steps, 1× PBS was used. The plates were blocked with 150 μl PBS-Casein for 2 hours at room temperature. Sera from mice were diluted 1:2 and incubated overnight at 4°C. On the following day, primary detection antibodies biotinylated anti-mouse CD81, anti-mouse gp100 (PMEL), and anti-mouse GPNMB were added at the concentration of 1 μg/mL. The plate was incubated for 1 hour at room temperature. The secondary detection antibody anti-rabbit IgG (H+L) HRP was added at the concentration 1:500, whereas Streptavidin/HRP was added at the concentration 1:1000. The details of antibodies are provided in Table 1. The plate was incubated for 1 hour at room temperature. The assay was developed with tetramethylbenzidine (TMB) and stopped with the equal volume of 1 M H2SO4 solution. Optical density (OD) measurements were taken at 450 nm using BioTek Cytation 5 cell imaging multimode reader.

### Animal model and vaccination experiment

All animal experiments were conducted in strict adherence to the Swiss Animal Act (455.109.1, September 2008, 5th edition), approved by the local ethics committee (#BE10/18, BE43/21 and BE30/24). All experiments were performed using female wild-type C57BL/6 mice aged 8–12 weeks, obtained from Harlan and housed in the central animal facility (CAF) at the Department of BioMedical Research, University of Bern. RAG1⁻/⁻ mice on a C57BL/6 background were bred and maintained under the same conditions at the CAF. The process of Qβ-VLPs expression, production, packaging and the generation of Qβ(CpG)-p33 for vaccination in B16F10 melanoma model was previously described^30^. For the experiments, three groups were established: Untreated, treated with non-specific Qβ(CpG)-VLPs, and treated with tumor-specific Qβ(CpG)-p33 VLPs. The vaccine was subcutaneously administered on days 3, 7, and 10 after the tumor transplantation at 50 μg/mouse diluted in 100µL PBS. The blood collection was performed on days 2, 9, and 14. The experiments were terminated 15 days after the tumor transplantation.

### Flow cytometry analysis of TILs

Tumors were digested in an enzyme mixture containing Collagenase D (1mg/mL in DMEM, Roche) for 30 minutes at 37 °C with agitation (200 rpm). Following the digestion tissue was passed through a 70 µm cell strainer. During processing, cells were repeatedly washed with complete DMEM using 50 mL Falcon tubes to remove debris. The resulting cell suspension was transferred to 15 mL tubes containing 2 mL of 35% Percoll and centrifuged at 1800 rpm for 25 min at room temperature to enrich for TILs. The pellet was resuspended in 200 µl of PBS supplemented with 0.1% BSA (flow cytometry buffer) and transferred to 96-well V-bottom plates for staining. Plates were centrifuged at 1200 rpm for 5 min, the supernatant was discarded, and cells were resuspended in flow cytometry buffer. The stained TILs were then passed through a cell strainer into 50 mL round-bottom tubes to remove residual tumor debris before acquisition. Samples were analyzed on Aurora 5L flow cytometer, and data were processed using GraphPad Prism (version 10.6.1). Tumor cell densities were calculated as the ratio between the total number of cells and the corresponding tumor weight (in mg). All antibodies (mAbs) were used at a 1:300 dilutions, while tetramers were used at a 1:50 dilution. The gating strategy sequentially excluded doublets and dead cells, followed by selection of lymphocytes and CD8⁺ T cells, which were further subdivided into p33⁺ populations based on tetramer staining.

### Human samples and ELISA

The study population consists of 5 melanoma patients and 5 healthy donors used as controls (not matched for age and sex). Patients were seen at the UPMC Hillman Cancer Center; data and samples were obtained from the University of Pittsburgh Melanoma SPORE Bank (IRB #970186). For HDs, data and samples were collected via IRB-approved protocol #991206. All study participants provided written informed consent. Blood samples were processed; plasma was separated, aliquoted into 2 mL cryotubes and stored at −80 °C until thawed and diluted 1:2 for analysis by ELISA. For these assays, the ELISA plates were coated with anti-human CD9, CD63, and CD81 antibodies (all at 1µg/mL, Thermo), whereas detection was done with anti-human gp100, and goat anti-mouse IgG HRP antibody (Thermo).

### Software and Statistical analysis

Illustrations were generated using BioRender. ChatGPT was used for language editing and formatting. Statistical analyses were performed using GraphPad Prism 9 (GraphPad Software). Statistical tests were applied as indicated in the corresponding figure legends. For comparisons between two groups, a two-tailed Mann–Whitney test was used. For comparisons involving more than two groups, the Kruskal–Wallis test with Dunn’s multiple comparisons test was applied. In vivo experiments were analyzed using two-way ANOVA with Tukey’s multiple comparisons test.

Pearson’s correlation analysis was used to assess associations between continuous variables and is reported as correlation coefficients (r) with two-tailed P values. Receiver operating characteristic (ROC) curve analysis was performed to evaluate biomarker discrimination, with performance quantified by the area under the curve (AUC) and 95% confidence intervals.

Unless otherwise indicated, a significance threshold of α = 0.05 was used. Statistical significance is indicated as P ≤ 0.05 (*), P ≤ 0.01 (**), P ≤ 0.001 (***), and P ≤ 0.0001 (****).

## Author contributions

PE conceptualized the study. PE and MM designed the experiments. All authors developed the methodology; EV isolation procedures were initially developed and supervised by SVG and DT; EV characterization techniques including LC-MS and ELISA were developed by PE and MS; Animal experiments were developed by PE, MM and MFB; Animal maintenance and ethical permission was managed by MM and ACV. Human patients and experiments were overseen by TW. Experiments were performed by MS, AG, ASC, JH, PE, and MM. Funding was acquired by PE and MM. MS, PE, and MM drafted the initial manuscript, before revisions and editing was done by all authors.

## Acknowledgements

We thank the Proteomics and Mass-Spectrometry Core Facility (PMSCF) of the University of Bern, led by Prof. Manfred Heller. We acknowledge Sandra Nansoz and Lee-Anne Brand for technical and administrative assistance. We thank Prof. Monique Vogel for scientific discussions.

## Funding

This study was supported by funding from the Holcim Foundation for Scientific Research, Novartis Foundation for Medical-Biological Research, the ACTERIA Foundation, and the Swiss National Science Foundation (SNSF), grant numbers 228774, 10001271 to PE, and Swiss Cancer Research, grant number KFS-5246-02-2021-R to MM, and SNSF grants 320030_219451 and 310030_219605, and the Swiss Cancer Research foundation (KFS-5936-08-2023-R) to SVG.

## Competing interests

MM and MFB have financial interests as shareholders in DeepVax GmbH, a spinoff from the University of Bern involved in the development of cancer immunotherapy.

## Data availability

All data are available on request

---

**Figure S1.**
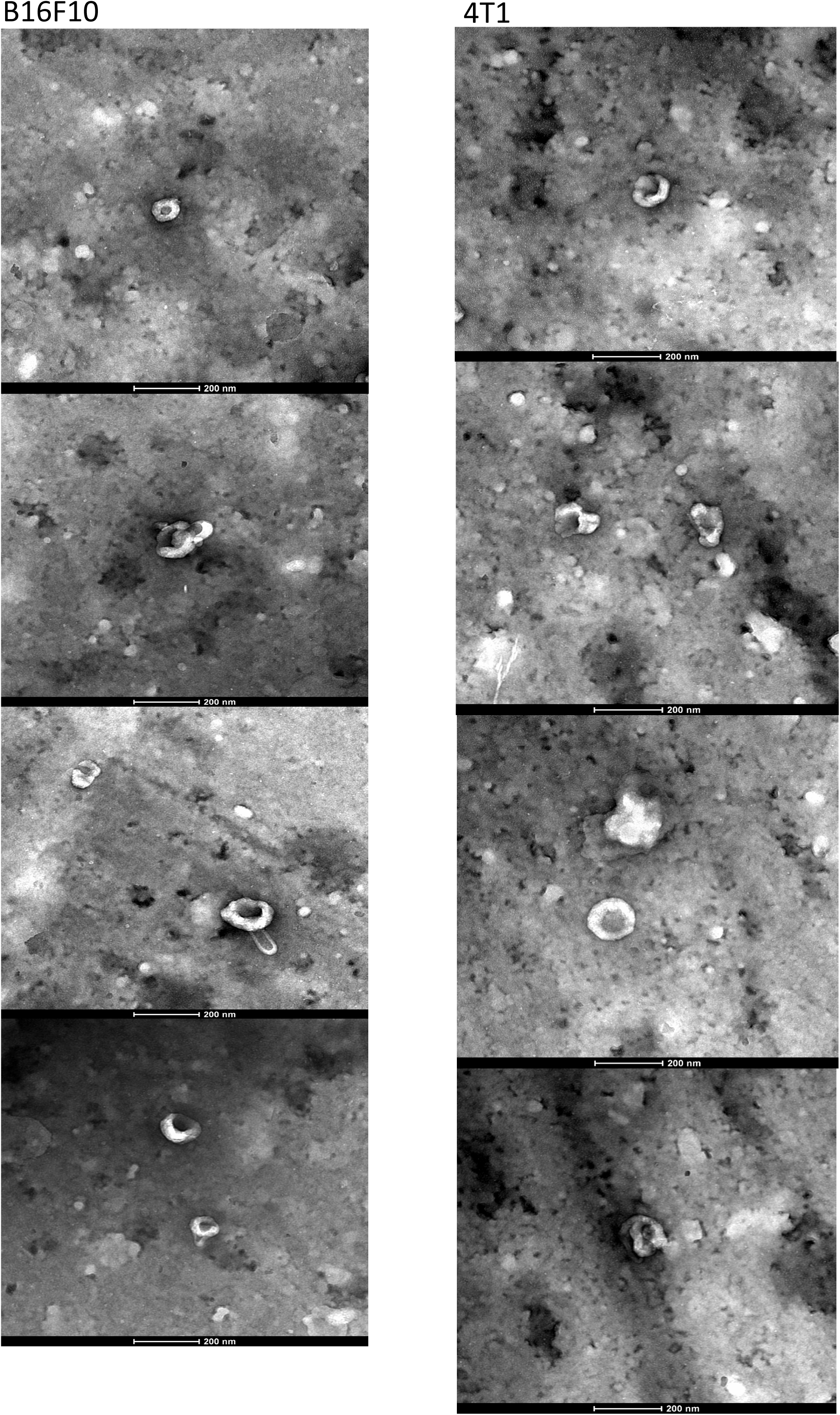
Additional TEM images of EV isolates for B16F10 (left panel) and 4T1 (right panel)

**Figure S2.**
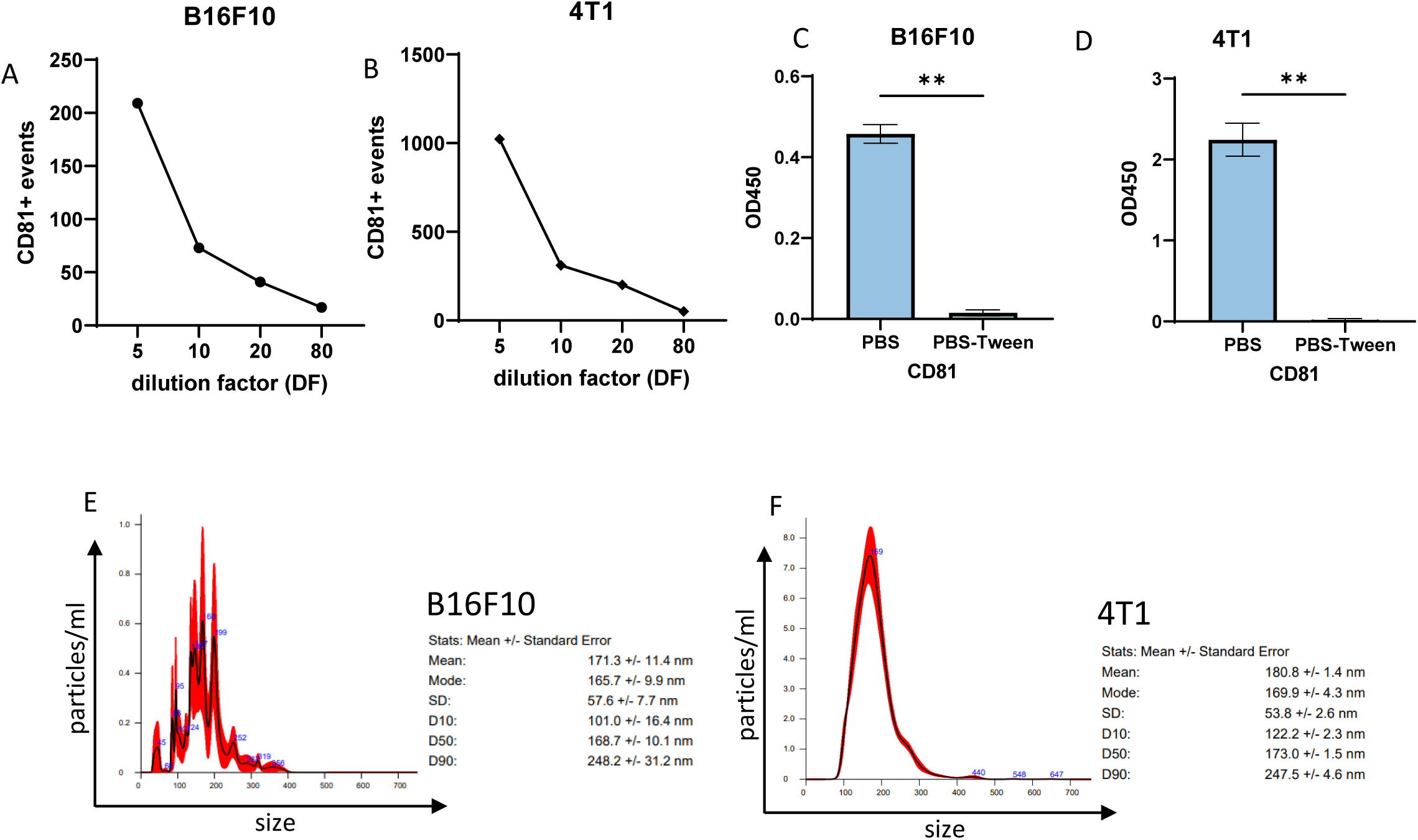
Supplementary ELISA data related to Fig. 1. Detection of CD81⁺ EVs upon serial dilution of isolated EVs from B16F10 (A) and 4T1 (B) cell lines. Effects of PBS–Tween (0.01%) treatment on CD81 signal in B16F10 (C) and 4T1 (D) EV preparations. Nanoparticle Tracking Analysis (NTA) of B16F10 (E) and 4T1 (F) EV preparations.

**Figure S3.**
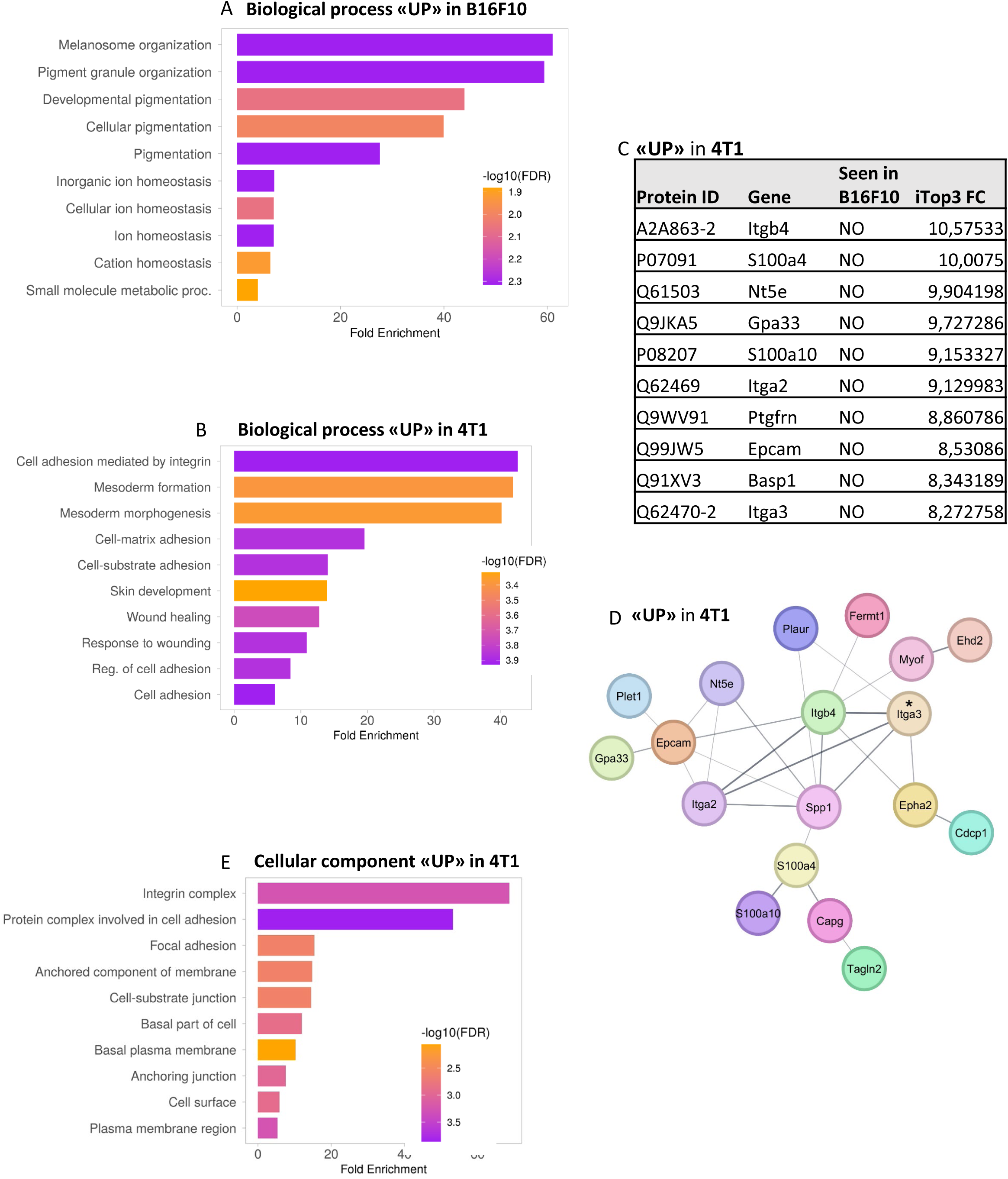
Supplementary LC–MS/MS analysis related to Fig. 2. GO biological process pathway enrichment for B16F10 (A) and 4T1 EVs (B), individual proteins enriched in 4T1 (C) and network representation (D), GO cellular component for B16F10.

**Figure S4.**
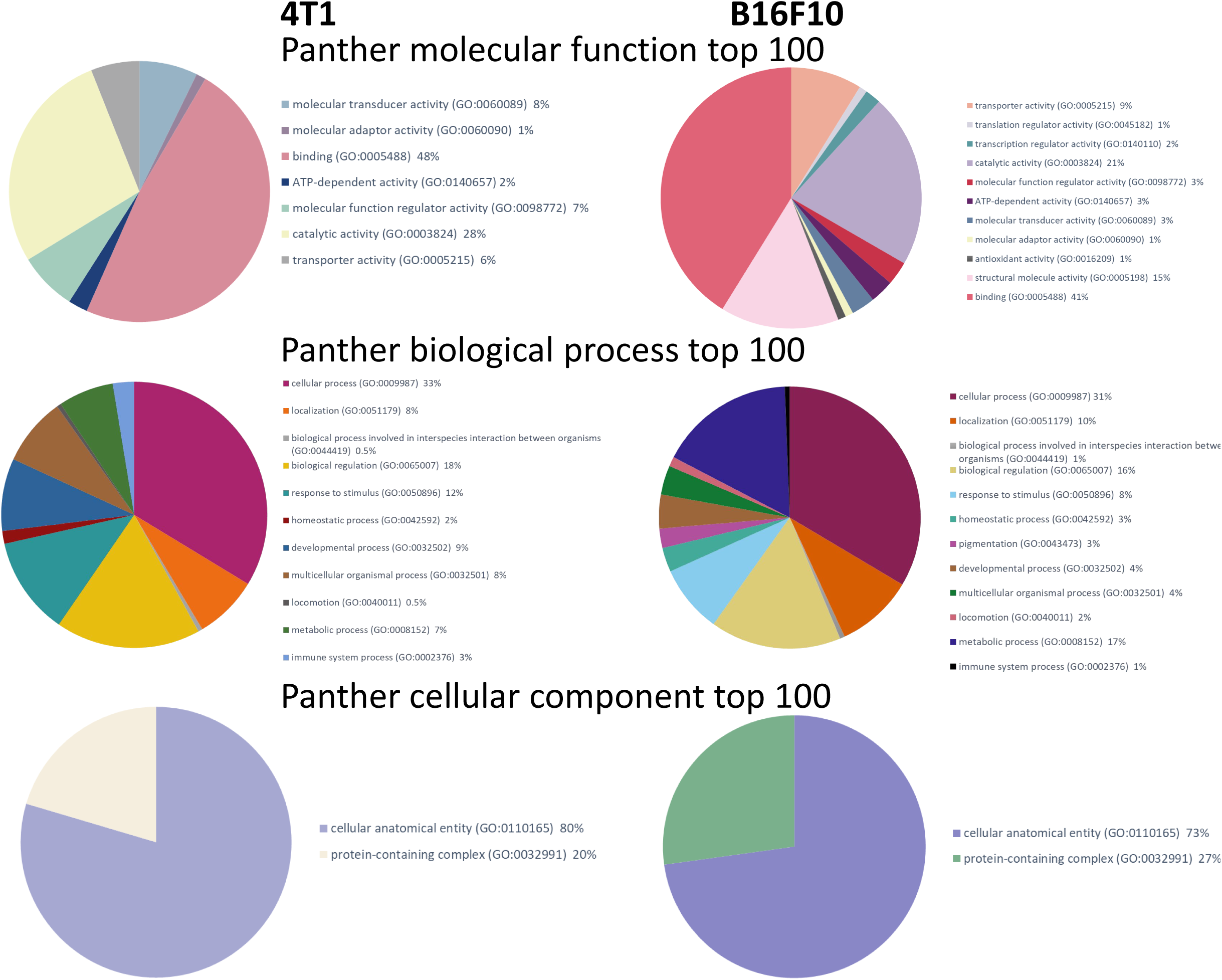
PANTHER-based pathway enrichment analysis as supplementary LC–MS/MS analysis related to Fig. 2. Top 100 enriched pathways across molecular function, biological process, and cellular component categories for B16F10 (right panels) and 4T1 (left panels) EVs.

**Figure S5.**
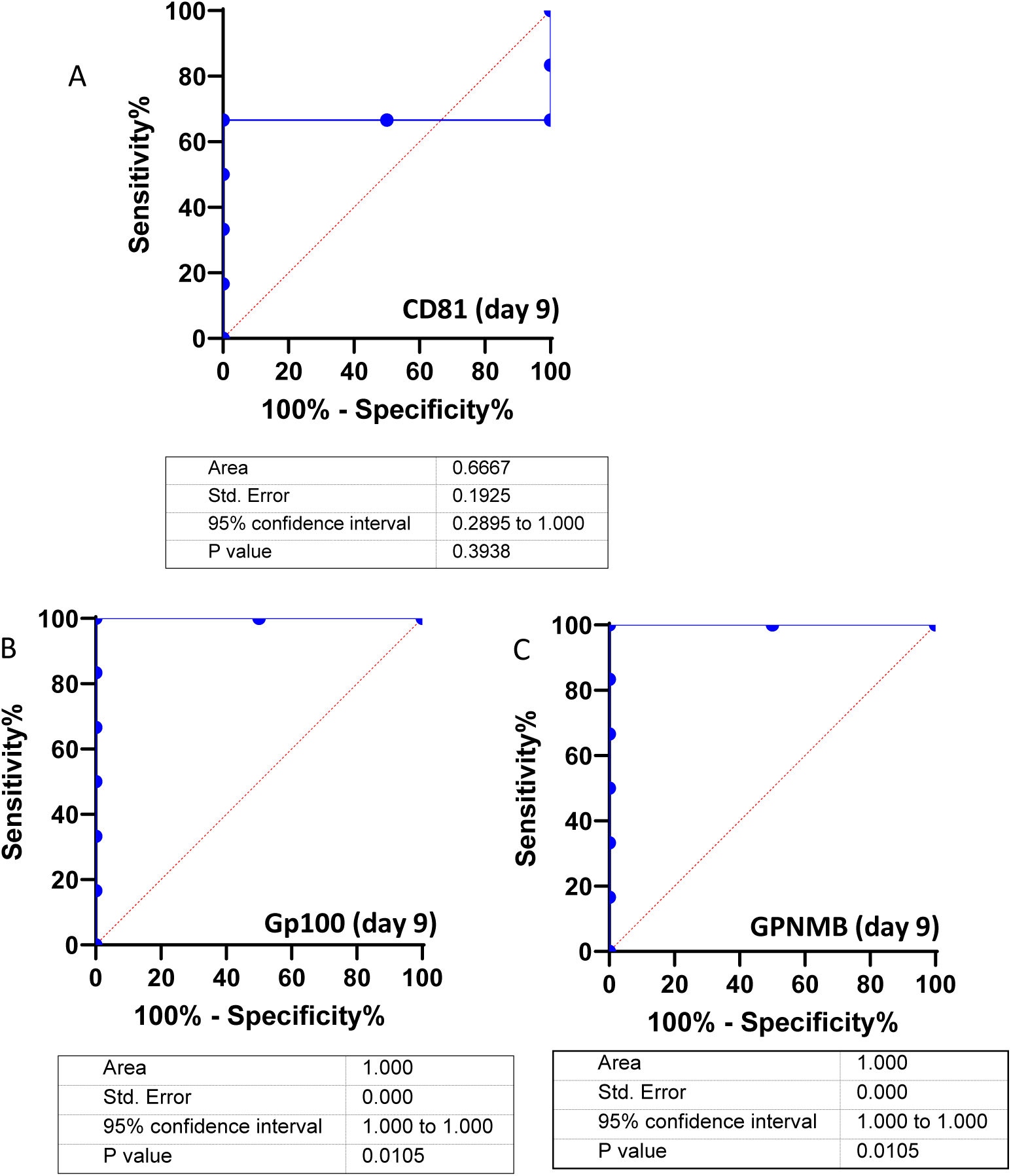
Supplementary analysis for Fig. 3. Receiver operating characteristic (ROC) curve analysis at day 9 for untreated versus treated animal for CD81⁺ EVs (A), gp100⁺ EVs (B) and GPNMB⁺ EVs (C).

**Figure S6.**
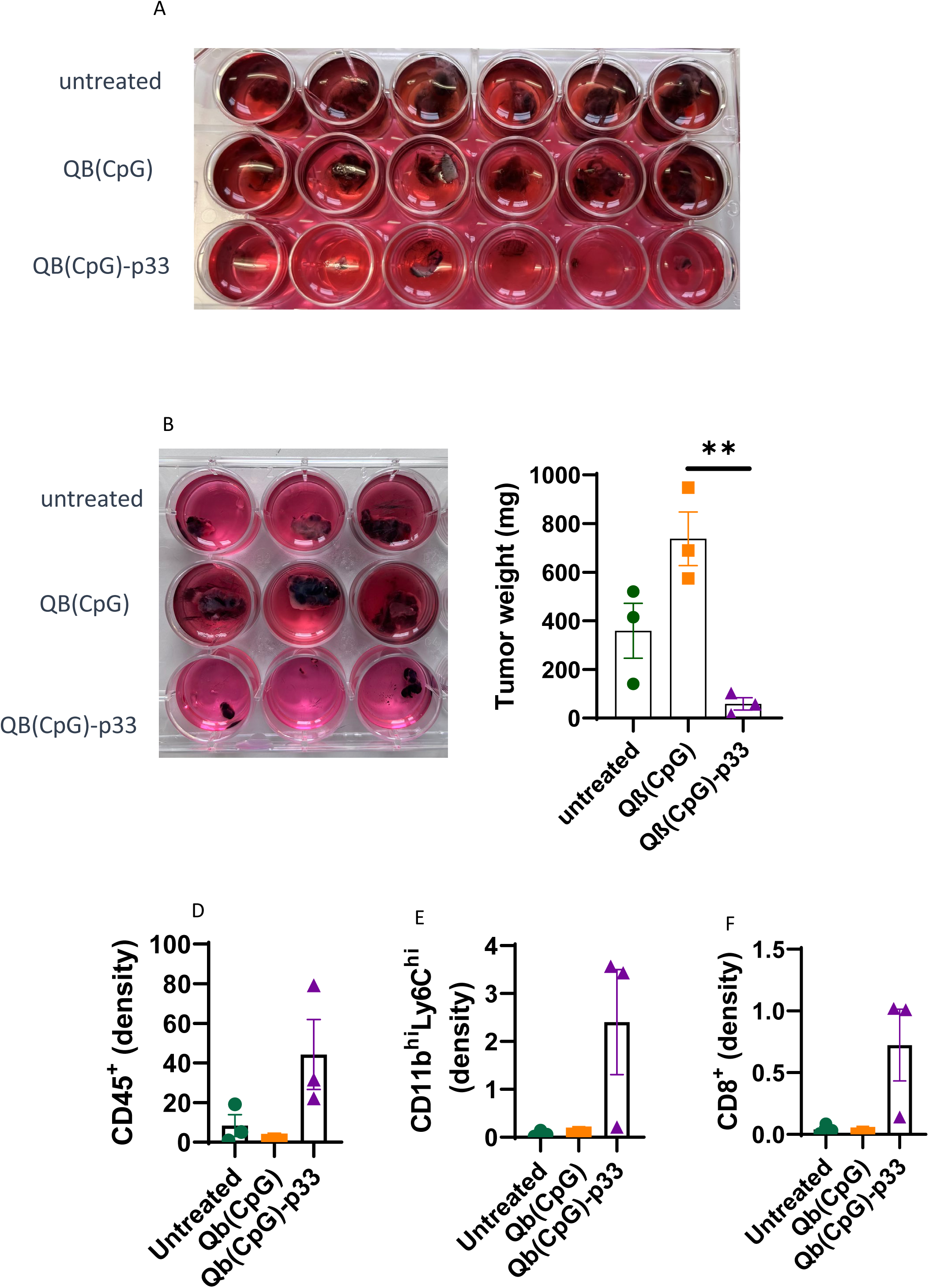
Supplementary data related to Fig. 2. Images of tumors and tumor weights from untreated, control-vaccinated Qβ(CpG), and antigen-specific Qβ(CpG)-p33 groups across two experiments (A, B) and tumor-infiltrating lymphocyte (TIL) densities (C, D, E) for experiment 2.

**Figure S7.**
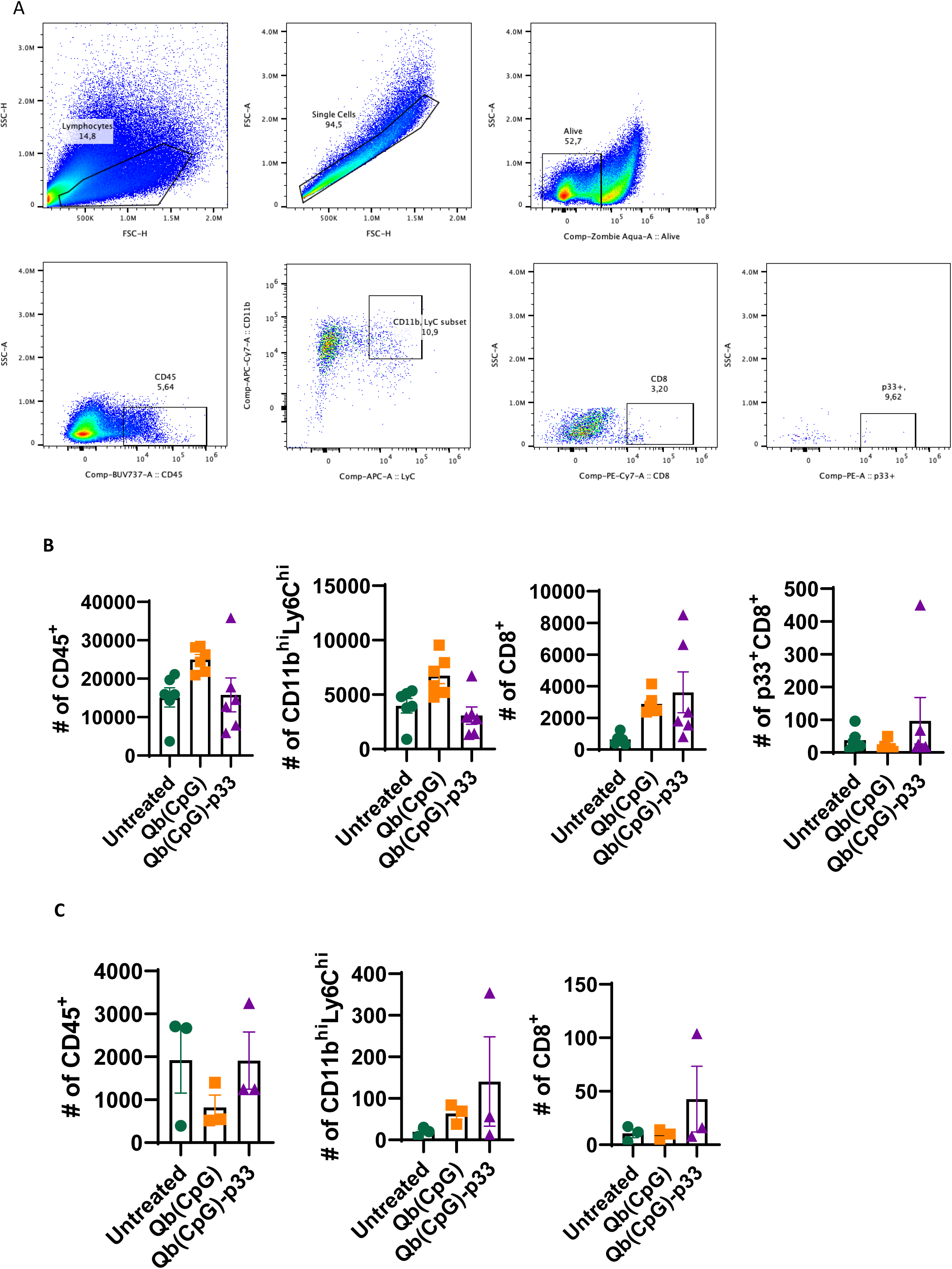
Supplementary data related to Fig. 3. Flow cytometry gating strategy for tumor-infiltrating lymphocytes, including CD45⁺ leukocytes, CD11b^hi^Ly6C^hi^ macrophages, CD8⁺ T cells, and p33 tetramer–specific CD8⁺ T cells (A). Absolute cell counts from two independent experiments are shown (B, C).

**Figure S8.**
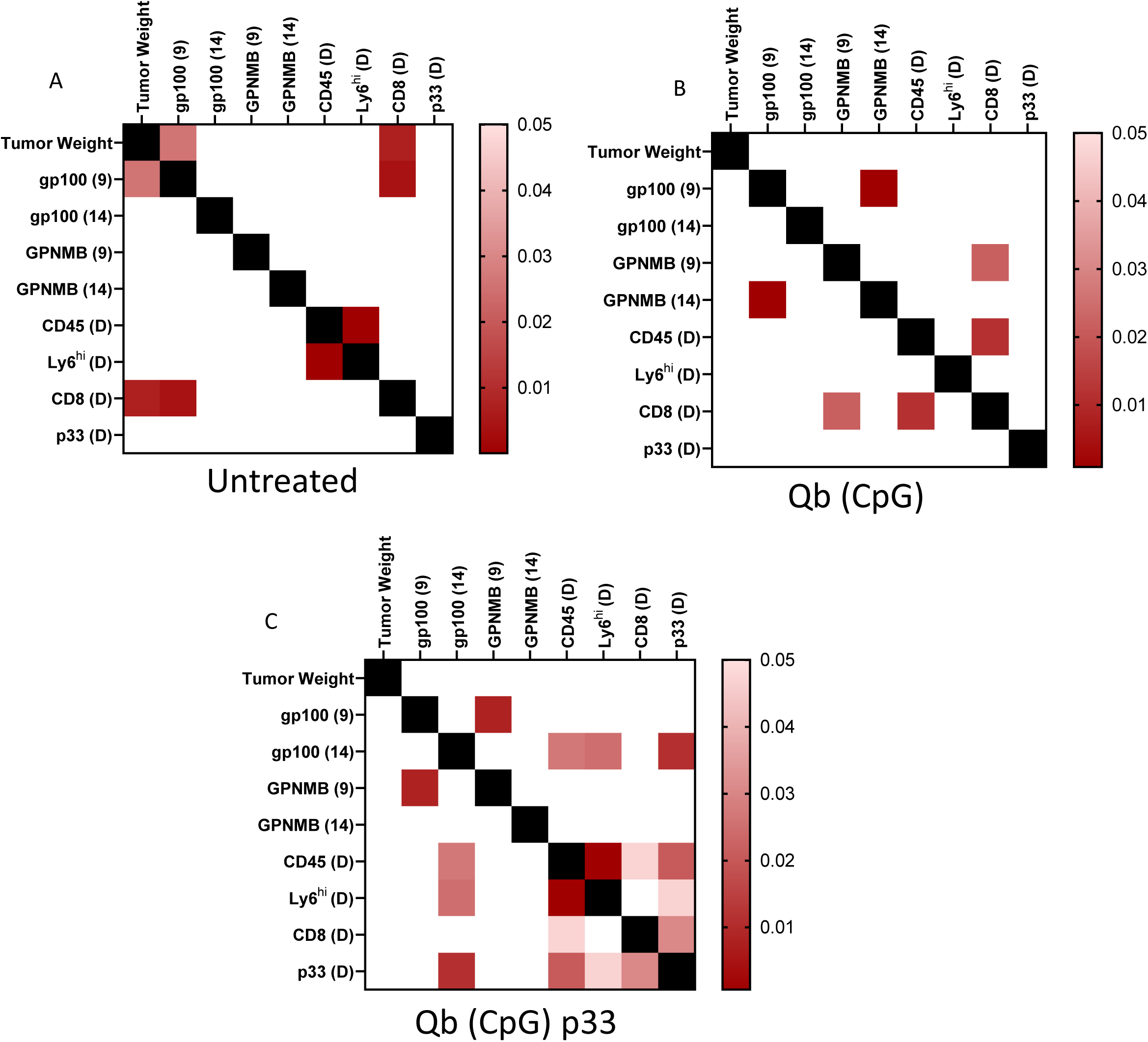
Supplementary data related to Fig. 3. Pearson correlation p-value matrices for untreated (A), control-vaccinated Qβ(CpG) (B), and antigen-specific Qβ(CpG)-p33 (C) groups.

**Figure S9.**
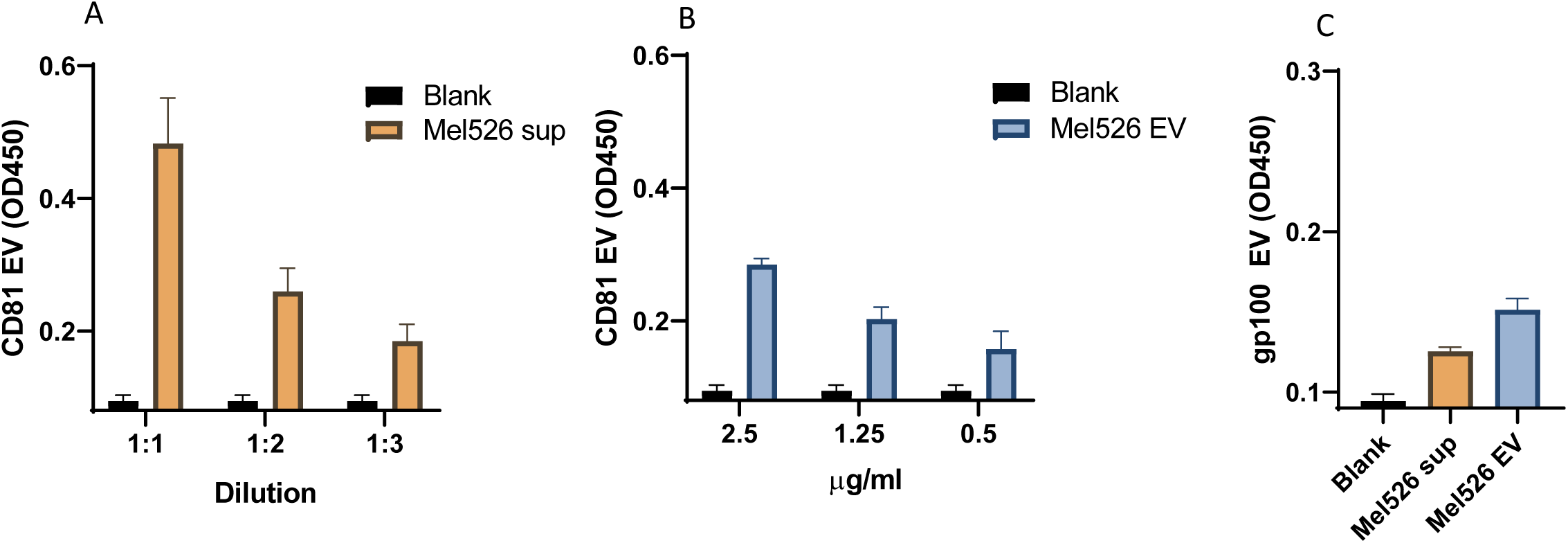
Supplementary data related to Fig. 4. CD81⁺ EV detection by ELISA in Mel526 cell supernatants (A) and isolated EVs (B), and gp100⁺ EV detection in supernatants and isolated EVs (C).

